# Fixing the Measurement Model Before the Selection Heuristics: Optimiser Repairs for Living-Cell Self-Driving Laboratories

**DOI:** 10.64898/2026.10.27.762131

**Authors:** Nura Brimo, Dilek Çökeliler Serdaroğlu

## Abstract

Closed-loop Bayesian optimisation assumes things that living cells do not provide. A measured cell is used up and cannot be measured again, cultures drift over a campaign, day-to-day batch offsets shift every reading, and cell-to-cell variation is heavy-tailed and mixed with probe failures. When an autonomous loop goes wrong under these conditions, the usual response is a selection-level repair: change the incumbent, replicate the current best, or explore more. The other option is to change what the model assumes about each measurement. We compared the two in a simulated living-cell laboratory with common random numbers, crossing two surrogate models with four selection repairs over three response surfaces, four pathology regimes and 60 seeds (8,820 campaigns of 30 rounds, five destroyed cells per round). A measurement-aware model, using a robust per-condition summary, shrunk per-condition noise and additive drift and day terms, lowered final simple regret against plain GP-EI by 0.042 [0.024, 0.061] under heavy tails and by 0.039 [0.013, 0.064] with all pathologies on, and raised 90% band coverage under drift from 0.83 to 0.95. It cost 0.015 [0.004, 0.027] on clean data. None of the selection-level repairs we tested helped either model once the data were contaminated, and using the best posterior mean as the incumbent raised regret in every regime for both models (+0.015 to +0.045). Because that variant is not integrated noisy expected improvement (NEI), we extended the study with an independent re-implementation of the simulator (10,260 further campaigns plus 4,320 for trajectory and sensitivity analyses). It reproduced the direction of the main effects with smaller magnitudes, showed that the measurement-aware gain persisted when drift was halved or doubled and when the probe-failure rate ranged from 5% to 20%, and found that integrated NEI did not hurt and gave the lowest regret when paired with the measurement-aware model. It also showed that offset-removed coverage overstates absolute calibration under drift. Within simulation, the measurement model is the first place to intervene; we have not tested real cell data.

## 1 Introduction

Self-driving laboratories have worked best where samples are stable and an experiment can be repeated on the same material [1–4]. Living cells are different. A cell probed with an optical trap, an AFM tip or a deformability channel is perturbed or lost, so a condition is only ever known through a fresh sample of cells. Stiffness distributions are broad [5], some readings are artifacts (a slipped bead, a detached cell), and the culture changes with passage and time in the incubator, on top of the usual batch effects [6].

When a loop converges on the wrong condition there are two places to intervene. Selection-level repairs change which condition is chosen next: using the posterior mean rather than the best noisy observation as the incumbent [7, 8], spending rounds on replicates of the current best, or exploring more [9]. Measurement-model repairs change what the surrogate assumes about the data: robust summaries of the destroyed cells [10, 11], condition-dependent noise [12], and explicit terms for time and batch [13]. Both are easy for an agent to add. We ask which one matters when the samples are alive.

### Contributions

(1) A simulator of living-cell closed loops with destructive measurement, drift, day effects, heteroscedastic heavy-tailed noise and probe failures, run with common random numbers so that every arm sees the same cells. (2) A comparison of two surrogate models crossed with four selection repairs, plus three ablations, over 8,820 campaigns. (3) Evidence, within this simulation, that the measurement model is where an agent should spend its effort first, and that the selection-level heuristics we tested do not help and one of them hurts. (4) An extension in an independent re-implementation that adds integrated NEI, regret trajectories, sensitivity to the size of the pathologies, and absolute calibration, which together mark where the main conclusion holds and where it needs qualification.

## 2 Related work

### Bayesian optimisation under noise

Expected improvement [14] needs an incumbent, which is ill-defined when observations are noisy. Plug-in fixes replace the best observation with the best posterior mean [8]; integrated NEI instead averages EI over the posterior of the latent function at the observed points [7], and is the default in BoTorch [15]. Reviews of the area [16, 17] treat noise mainly as a property of the acquisition rule. We treat it as a property of the measurement and ask how much of the problem can be solved before the acquisition step.

### Robust and heteroscedastic surrogates

Student-*t* and outlier-aware GPs make BO robust to corrupted readings [11], and input-dependent noise models capture conditions that are intrinsically more variable [12]. Our measurement-aware model uses simpler ingredients: a Huber location per condition [10] and empirical-Bayes shrinkage of per-condition variances [18]. These are cheap and transparent to a human supervisor, and they exploit the fact that each round yields several cells at one condition.

### Time-varying optimisation

Drift has been handled by contextual and time-varying GP bandits [13, 19]. We use additive time and day kernels, which suits an objective whose level shifts while its shape is stable.

### Self-driving laboratories

Most demonstrations are in chemistry and materials [1–4], where samples can be re-measured. Living-cell loops lack that property, and we know of no controlled comparison of model-level and selection-level repairs for this setting.

## 3 Setup

### Simulated laboratory

The design space is *x∈* [0, 1]^2^ (for example a drug dose and an osmotic load). A campaign runs *T* = 30 rounds: six random initial conditions, then 24 chosen by the optimiser. In round *t* the laboratory measures *n* = 5 fresh cells at the chosen *x*, and those cells are never measured again. Cell *j* reads

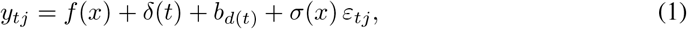

where *f* is the population response scaled to [0, 1] on the grid, *δ*(*t*) is a linear culture drift reaching 0.35 by the last round, *b*_*d*_*∼ N* (0, 0.1^2^) is an offset for each four-round day, *σ*(*x*) = 0.25 (1 + 1.2*x*_1_), and *ε* is unit-variance Student-*t*_3_ noise. With probability 0.1 a reading is replaced by a probe failure, *f* (*x*) + *δ*(*t*) + *b*_*d*_ *± U* (1, 2). Four regimes switch these effects on in blocks: clean (Gaussian noise, *σ* = 0.25, no drift), drift (drift and day offsets, Gaussian noise), tails (heteroscedastic *t*_3_ noise and failures) and full (everything). The three surfaces are a narrow global peak beside a broad local one, a negated Branin function, and a sigmoidal dose response with an optimal window, a toxicity dip and a small linear trend in *x*_2_. For a given seed, every arm sees the same initial design, drift, day offsets and noise draws.

### Surrogate models

Both models are Gaussian processes (GPs) with a squared-exponential kernel and hyperparameters chosen by marginal likelihood on a grid. The plain model fits the sample mean of each condition with a learned homoscedastic noise. The measurement-aware model changes three things. It summarises each condition by a Huber location (*k* = 1.345) [10], scaled by that condition’s median absolute deviation (MAD) shrunk toward the median MAD across conditions with eight prior degrees of freedom [18]. It passes the shrunk variance, divided by *n* and inflated by 1.1 for the Huber estimator, to the GP as fixed per-point noise, plus a learned nugget. And it adds a smooth kernel over time and a same-day kernel [13], which are dropped when predicting *f* . Three ablations remove the time and day kernels, the Huber summary and shrinkage (sample mean with its raw variance instead), or the per-condition noise (homoscedastic instead).

### Acquisition and selection repairs

The default rule is EI [14] over a 32 *×* 32 grid, with the best observed summary as the incumbent. The selection repairs, applied to either model, are: a posterior-mean incumbent (the best posterior mean at the sampled points, labelled “noisy-EI incumbent” in Figure 1 and Table 1); a replicate incumbent (every fourth round re-measures the current recommendation with new cells); UCB (*µ* + 2*σ* in place of EI); and all selection repairs (posterior-mean incumbent plus replication). The posterior-mean incumbent is a plug-in approximation. It is not the integrated NEI of Letham et al. [7], which we add in Section 5.

**Table 1.**
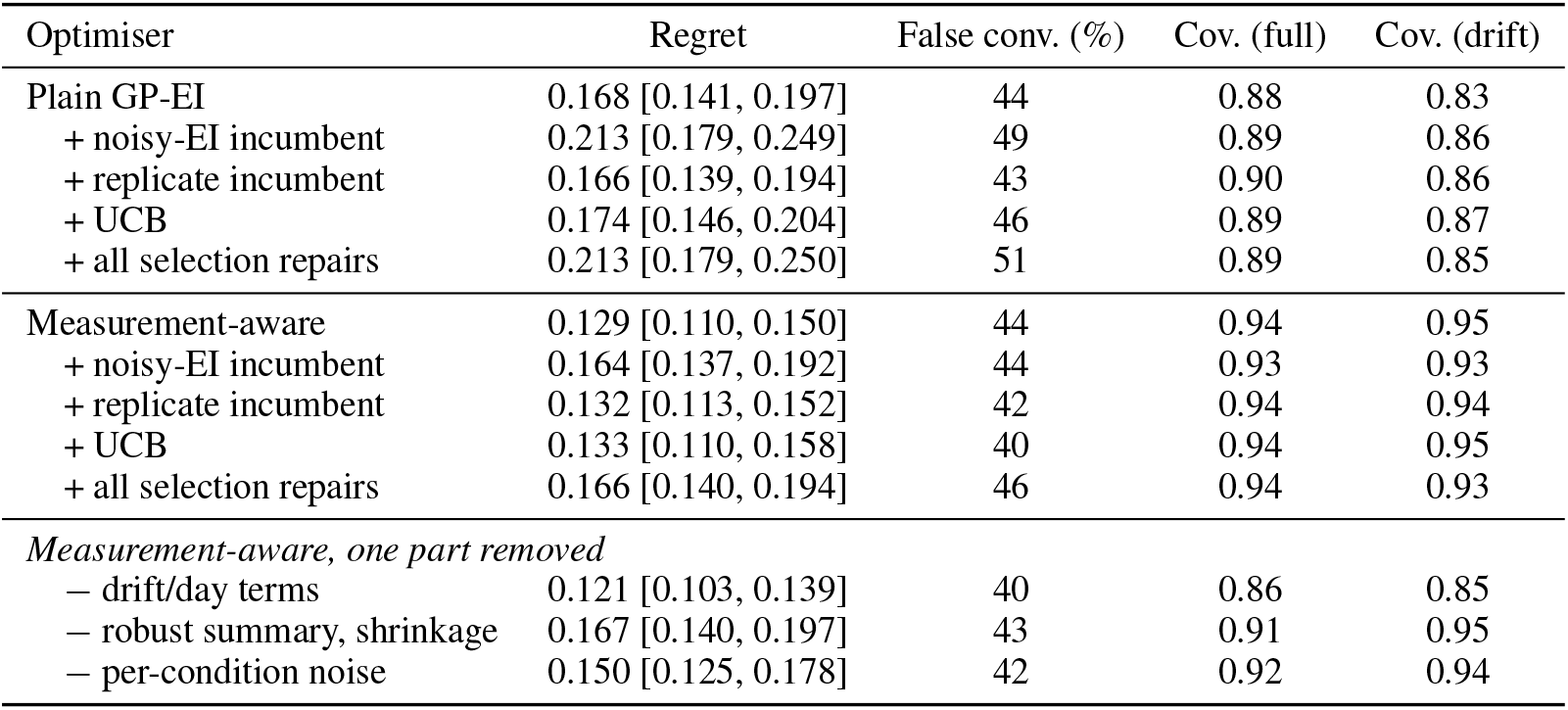
Full regime (all pathologies), 180 campaigns per row. Regret is the mean [95% bootstrap interval]; false convergence is the share of campaigns with regret *>* 0.1; coverage is for the nominal 90% band of *f* (offset removed). The last column is coverage in the drift regime.

**Figure 1.**
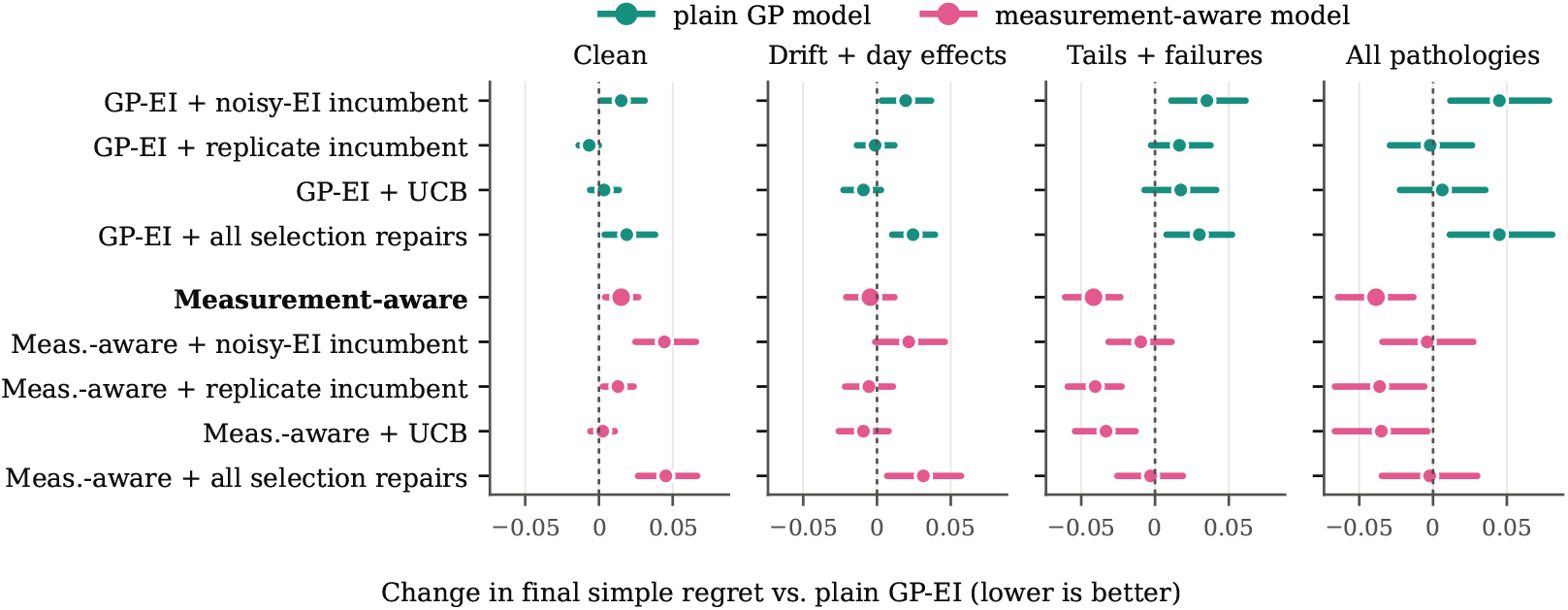
Paired change in final simple regret relative to plain GP-EI, pooled over three surfaces and 60 seeds per regime (mean and 95% bootstrap interval; objective range is 1). Points left of zero are better. “Noisy-EI incumbent” denotes the posterior-mean-incumbent variant, not integrated NEI.

### Metrics

After round 30 each arm recommends the grid point with the highest posterior mean of *f* . We report simple regret 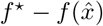 on the [0, 1] scale; false convergence, the share of campaigns with regret above 0.1; and calibration, the coverage of the nominal 90% posterior band for *f* over the 1,024 grid points after removing a constant offset (the absolute level is not identifiable under drift, so this measures the shape and width of the band, not its absolute position; Section 5.5 reports absolute coverage). Every model and selection combination ran in all four regimes (180 campaigns each: 3 surfaces *×* 60 seeds); the ablations ran in the three contaminated regimes. Intervals are 95% bootstrap intervals, and differences between arms are paired by seed.

## 4 Results

### The measurement model helped where the data were contaminated

Against plain GP-EI, the measurement-aware model lowered regret by 0.042 [0.024, 0.061] under heavy tails and by 0.039 [0.013, 0.064] with all pathologies on (Figure 1). It was neutral under drift alone ( *−*0.004 [*−* 0.021, 0.012]) and cost 0.015 [0.004, 0.027] on clean data, which is the price of a robust summary and shrunk noise when the data do not need them. With all pathologies on it lowered mean regret on every surface: 0.237 vs. 0.302 on the two-peak surface, 0.036 vs. 0.046 on Branin and 0.116 vs. 0.157 on the dose response. Against the optimiser a practitioner might build after seeing failures, plain GP-EI with all selection repairs, the gap was 0.084 [0.056, 0.113].

### Selection repairs did not help, and one hurt

The posterior-mean incumbent raised regret in every regime for both models: +0.015, +0.019, +0.035 and +0.045 for the plain model (clean, drift, tails, full) and +0.029, +0.026, +0.032 and +0.035 for the measurement-aware model, with every interval above zero. A likely reason is that with five cells per condition the posterior mean over-credits the region that has been sampled most, and anchoring the incumbent there keeps EI from leaving it; we did not test this explanation directly. Replication changed regret by *−*0.002 to +0.002 on the measurement-aware model and gave the plain model at most a marginal gain on clean data ( *−*0.007 [ *−*0.014, 0.000]). UCB helped the measurement-aware model on clean data ( *−*0.012 [ *−*0.022, *−*0.003]) and nowhere else. Once the data were contaminated, no selection repair improved on either model’s default.

### Different parts of the model do different jobs

The ablations separate regret from calibration (Table 1, Figure 2). The robust summary carried most of the regret gain: without it, regret rose by 0.038 [0.011, 0.066] under all pathologies and 0.042 [0.022, 0.063] under heavy tails. Per-condition noise added a smaller, uncertain gain (+0.020 [ *−*0.003, 0.044] when removed). The time and day kernels did not change regret ( *−*0.009 [ *−*0.026, 0.008] when removed; the point estimate is slightly in favour of dropping them), but they carried the calibration. Without them, coverage fell from 0.95 to 0.85 under drift and from 0.94 to 0.86 under all pathologies. Plain GP-EI covered only 0.83 under drift, because it has no term for time and credits the drift to *x*. Not every ablation favours the full model, so the regret gain should be attributed mainly to the robust summary, and the calibration gain to the time and day terms.

**Figure 2.**
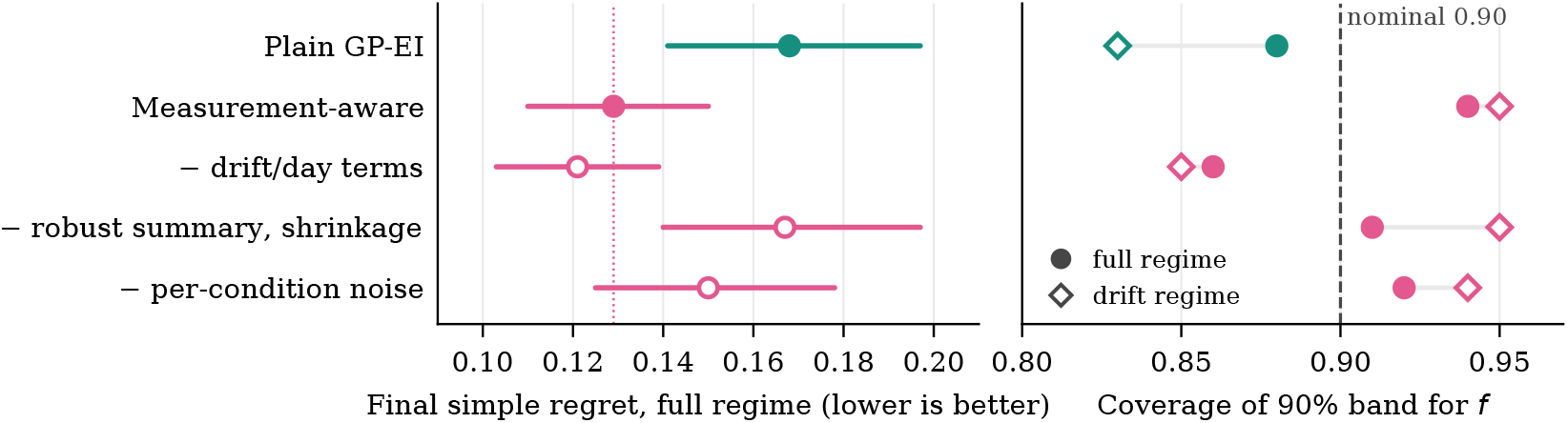
The ablations separate regret from calibration (data from Table 1). Left: final simple regret in the full regime (mean and 95% bootstrap interval); the dotted line marks the full measurement-aware model. Right: coverage of the nominal 90% band of *f* in the full regime (filled circles) and the drift regime (open diamonds). Removing the robust summary costs regret; removing the time and day terms costs calibration.

### Some failures are budget failures

False convergence stayed near 44% for both default models under all pathologies. Most of it came from the two-peak surface (83 to 92% of campaigns across arms), where 30 rounds of five cells rarely find a narrow global optimum beside a broad decoy, and from the dose response (27 to 52%). On Branin every arm stayed below 0.06 mean regret. No repair we tested fixes a budget that is too small for the surface.

## 5 Extended experiments

The meta-review raised three points that the original experiments could not answer: the acquisition comparison did not include integrated NEI, the noise and drift settings were our own choice, and calibration was measured only after offset removal. We addressed them in an independent re-implementation of the simulator, written from the specification in Section 3. Details that Section 3 does not fix (exact surface formulas, hyperparameter grids, and the incumbent of the measurement-aware model, here the best drift-corrected summary) were set independently. The results below are therefore a separate experiment, not a replication of Table 1, and we compare them to the main study only in direction.

### 5.1 What carried over

The re-implementation ran every arm of the main study plus integrated NEI for both models (10,260 campaigns, same design and seeds per arm). It reproduced the direction of the main effects but not their size. Plain GP-EI reached a mean regret of 0.194 [0.164, 0.224] in the full regime (0.168 in Table 1) and the measurement-aware model 0.170 [0.147, 0.193] (0.129). The paired gain of the measurement-aware model was *−* 0.021 [ *−*0.038, *−*0.002] under heavy tails and *−*0.024 [ *−*0.045, *−*0.003] with all pathologies, against *−*0.042 and *−*0.039 in the main study. Offset-removed coverage under drift rose from 0.81 to 0.89 (0.83 to 0.95), and removing the time and day terms lowered it from 0.89 to 0.80, so the calibration role of those terms carried over. Two findings did not. Removing the robust summary changed regret by only +0.004 [ *−* 0.012, 0.020], so the attribution of the regret gain to that component is specific to the original implementation. And the posterior-mean incumbent, although it raised regret under heavy tails and the full regime for the plain model (about +0.023) and under drift for the measurement-aware model (+0.028 [0.014, 0.042]), helped the plain model on clean data and was neutral elsewhere.

### 5.2 Integrated noisy EI

For each of 24 posterior draws of the latent function at the observed points, candidates are conditioned on that draw without noise and scored by EI against the draw’s best value; the scores are averaged [7]. Integrated NEI did not hurt either model (Table 2). With the measurement-aware model it reached a full-regime regret of 0.155 [0.135, 0.174], the lowest of all arms, against 0.170 for EI; the paired difference was *−*0.016 [ *−*0.034, 0.002] in the full regime and *−*0.020 [ *−*0.035, *−*0.005] under heavy tails. With the plain model it made no reliable difference. The two repairs therefore compose rather than compete: the measurement model supplies a cleaner posterior, and integrated NEI uses it better. This is why we no longer claim that the acquisition rule should be left alone; the claim we keep is about the selection heuristics in Section 4.

**Table 2.** Re-implementation. Paired change in final simple regret by regime (mean [95% bootstrap interval], 180 pairs; negative is better). NEI is integrated noisy EI; PM is the posterior-mean incumbent; EI is the default rule.

| Comparison | Clean | Drift | Tails | Full |
| --- | --- | --- | --- | --- |
| Aware vs. plain (both EI) | −0.000<br>[−0.015, 0.014] | −0.015<br>[−0.028, −0.003] | −0.021<br>[−0.038, −0.002] | −0.024<br>[−0.045, −0.003] |
| Plain: NEI vs. EI | −0.002<br>[−0.017, 0.010] | −0.008<br>[−0.019, 0.004] | −0.006<br>[−0.022, 0.009] | +0.003<br>[−0.017, 0.022] |
| Aware: NEI vs. EI | −0.010<br>[−0.019, 0.000] | −0.005<br>[−0.015, 0.004] | −0.020<br>[−0.035, −0.005] | −0.016<br>[−0.034, 0.002] |
| Plain: PM vs. EI | −0.013<br>[−0.028, −0.001] | +0.001<br>[−0.012, 0.014] | +0.023<br>[0.003, 0.043] | +0.023<br>[0.002, 0.045] |
| Aware: PM vs. EI | +0.007<br>[−0.002, 0.017] | +0.028<br>[0.014, 0.042] | +0.003<br>[−0.014, 0.020] | +0.017<br>[−0.000, 0.034] |
| Aware + NEI vs. plain + EI | −0.010<br>[−0.024, 0.001] | −0.021<br>[−0.033, −0.009] | −0.040<br>[−0.057, −0.024] | −0.039<br>[−0.059, −0.020] |

### 5.3 When the gain appears

Figure 3 tracks the recommendation through the campaign. All arms start from the same initial design (regret 0.29 after round 6). The measurement-aware model pulls ahead within six acquisition rounds: by round 12 its regret is 0.203 against 0.247 for plain GP-EI, a paired gap of *−*0.044 that narrows to *−*0.024 by round 30 as the plain model catches up. The measurement-aware model with EI flattens after round 24, while integrated NEI keeps improving (0.164 to 0.155). The posterior-mean incumbent tracks plain GP-EI until about round 12 and then falls behind, consistent with an incumbent that anchors on the most-sampled region. For a laboratory whose budget is shorter than 30 rounds, the measurement model matters more than the final-round numbers suggest.

**Figure 3.**
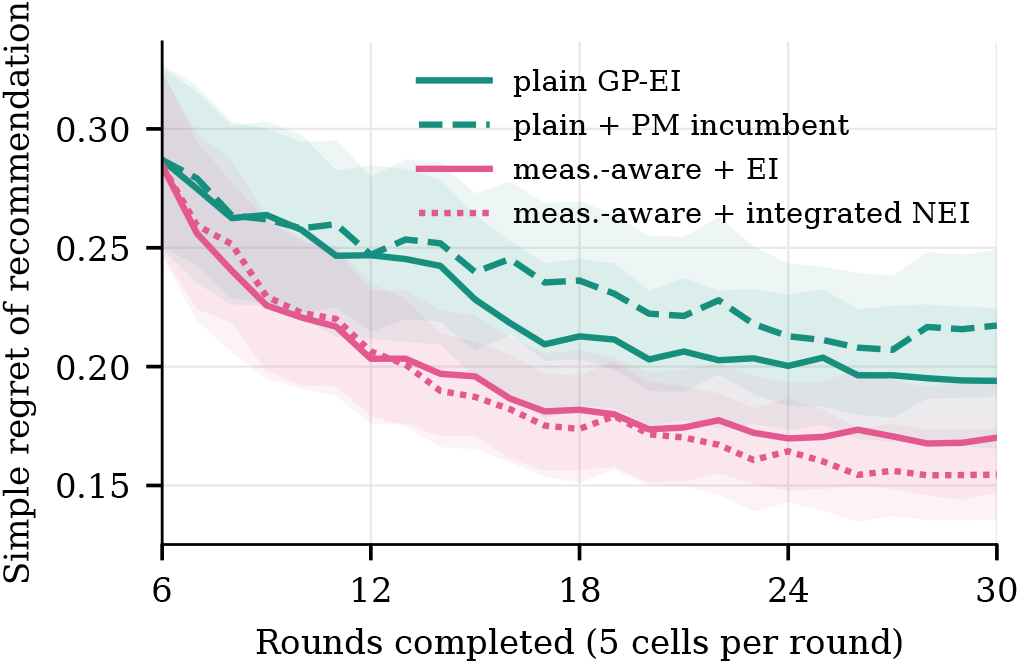
Re-implementation, full regime. Simple regret of the current recommendation after each round (mean over 180 campaigns; shaded 95% bootstrap band). Round 6 is the end of the random initial design.

### 5.4 Sensitivity to the pathology settings

The drift and failure parameters were our own choices, so we varied them one at a time in the full regime (Figure 4). With drift halved to 0.175 or doubled to 0.7, the measurement-aware model lowered regret by 0.036 [0.016, 0.056] and 0.046 [0.024, 0.069]; with the failure rate at 5% or 20%, by 0.028 [0.012, 0.046] and 0.027 [0.001, 0.053]. Every interval stayed below zero. Adding integrated NEI increased the gain in every setting (*−*0.039 to *−*0.064), and the largest gain came with the largest drift. The posterior-mean incumbent had a positive point estimate in all five settings (+0.006 to +0.023), but its interval excluded zero only at the default. Within this range, the ordering of the three arms did not depend on the settings we chose.

**Figure 4.**
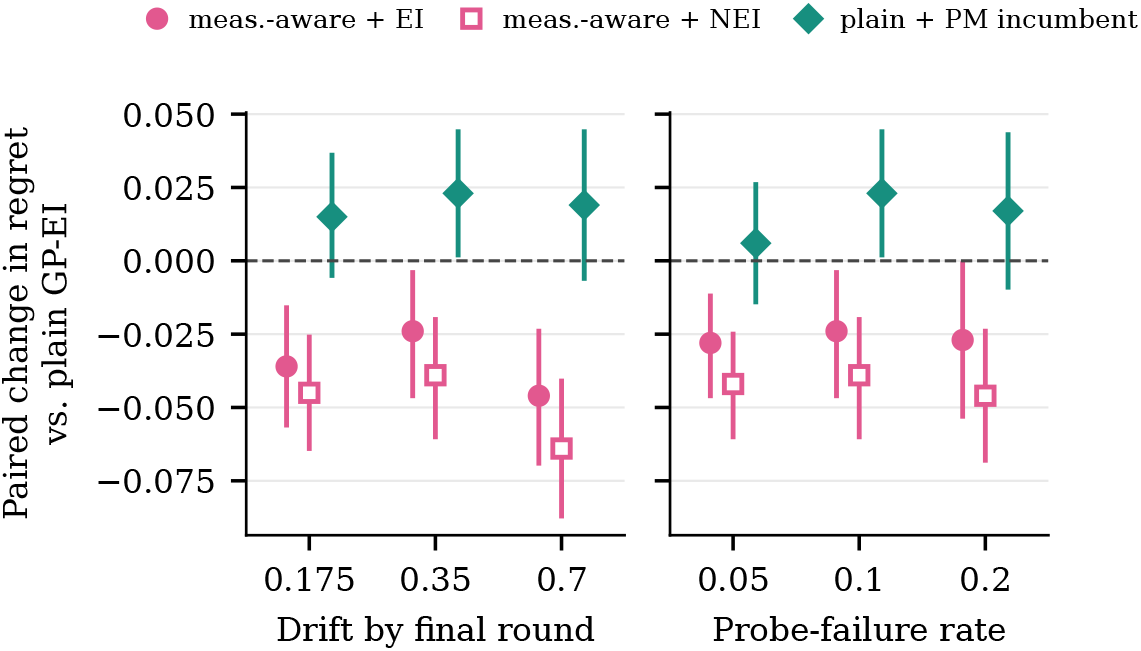
Re-implementation, full regime. Paired change in final regret against plain GP-EI (mean and 95% interval, 180 pairs) as drift (left; failure rate 0.1) and probe-failure rate (right; drift 0.35) vary. The middle setting in each panel is the default.

### 5.5 Absolute calibration

The meta-review noted that calibration was assessed only after offset removal. Table 3 shows why that matters. Without the offset correction, coverage fell for every model, and it fell furthest for the measurement-aware model (0.89 to 0.42 under drift), which had the best offset-removed coverage. The measurement-aware model gets the shape and width of *f* right, but when it separates *f* from the time and day terms it cannot tell which constant belongs to which, because the data contain no reference for the absolute level. The plain model is probably less wrong in level because it partly folds the drift into *f*, which is also what costs it shape calibration. An agent that needs absolute values, for example to compare a stiffness against a clinical threshold, should therefore anchor the level directly, for instance by measuring a fixed control condition each day. We did not simulate this design.

**Table 3.**
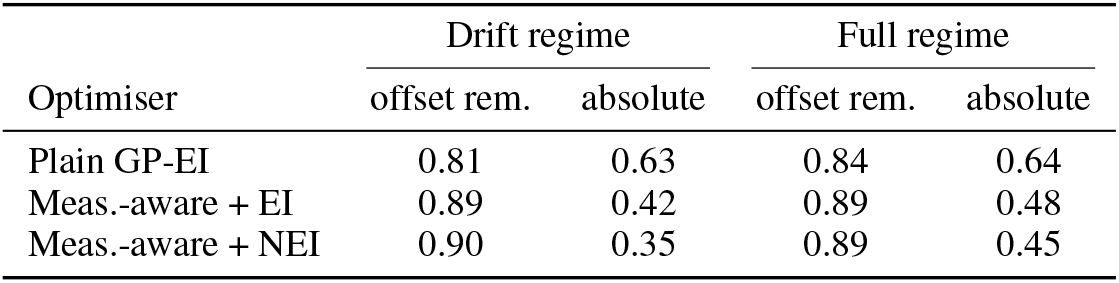
Re-implementation. Coverage of the nominal 90% band of *f* with and without removing a constant offset.

## 6 Discussion

For an agent running a living-cell loop, these results suggest an order of work. First, get the measurement model right: summarise destroyed cells robustly, give each condition its own shrunk noise, and model time and day explicitly. Second, check calibration, because the time and day terms are what make the posterior shape honest under drift, and calibration is what a human supervisor needs to decide whether to trust a recommendation; if absolute values matter, add a daily control. Third, treat plug-in selection heuristics with suspicion. In the original simulator the posterior-mean incumbent was harmful in every setting we ran, and replication and UCB gave no reliable benefit on contaminated data. Principled noise-aware acquisition is a different matter: integrated NEI did not hurt in the re-implementation and helped once the measurement model was in place, so it is a reasonable second step after the first. Because the measurement-aware model costs a little on clean data, an agent should switch it on from evidence, such as a replicate spread too wide for a Gaussian model or a control condition that moves from day to day, rather than by default.

### Limitations

This is a simulation. Drift was additive and linear, the design space was two-dimensional, each round measured one condition, and acquisition was optimised on a grid with a fixed kernel family. Real cultures can drift in ways that interact with treatment, and real probe failures are rarely symmetric. The noise and drift parameters were chosen by us and not fitted to measurements; the sensitivity analysis covers a factor of two in drift and four in failure rate, not other noise shapes. Integrated NEI, the trajectories, the sensitivity analysis and absolute calibration were evaluated only in the re-implementation, which reproduced the direction but not the size of the main effects and did not reproduce the attribution of the regret gain to the robust summary. The main gap is the absence of real cell data; the next step is to run the same arms on recorded optical-tweezers and AFM measurements from living cells, replaying them in silico before any closed-loop deployment.

## 7 Conclusion

In simulated living-cell campaigns, changing what the model assumes about each measurement improved both regret and calibration, while the plug-in selection heuristics an agent might reach for did not. Integrated NEI is compatible with this conclusion and adds to the gain. Absolute calibration under drift needs an experimental anchor that no surrogate can supply on its own.

## Code availability

Code for both simulators and all arms will be released publicly.

